# Transglutaminase 2 function in glioblastoma tumor efferocytosis

**DOI:** 10.1101/2024.08.29.610293

**Authors:** Margarita Lui, Filiz Sevinc, Mara Elgafarawi, David G. Munoz, Jeffrey W Keillor, John Sinclair, Dragosh Catana, John Woulfe, Ian AJ Lorimer

**Author notes:** Corresponding author; Ian Lorimer, The Ottawa Hospital Cancer Centre, 3rd Floor, 501 Smyth Road, Ottawa, ON, K1H 8L6.

## Abstract

Glioblastoma is an aggressive and incurable type of brain cancer. Regions of tissue necrosis are a distinctive pathological feature of this cancer. These arise through thrombosis of tumor vasculature, driven by tumor-derived pro-coagulation factors. In studies of transglutaminase 2 (TGM2), we observed that TGM2 mRNA expression in glioblastoma was primarily in a subset of tumor-infiltrating myeloid cells with hypoxia gene expression signatures. Analysis of xenograft and human glioblastoma samples by immunohistochemistry showed that macrophages in the vicinity of necrotic regions expressed very high levels of TGM2. These macrophages were engaged in the phagocytosis of apoptotic cells, a process known as efferocytosis. In cell culture, incubation of macrophages with apoptotic cells induced TGM2 expression in macrophages, and TGM2 inhibitors blocked efferocytosis. In patient-derived glioblastoma organoids cultured in 5% O_2_, a basal level of apoptosis was observed, and endogenous macrophages were observed in the process of clearing apoptotic cells. Clearance of apoptotic cells was reduced in organoids treated with a TGM2 inhibitor. Apoptotic cells and efferocytosis were both markedly lower in organoids grown in 20% O_2_. These data, together with previous work, define a model in which necrotic regions in glioblastoma induce hypoxia-driven apoptosis, which in turn promotes efferocytosis by macrophages. TGM2 is both a marker of efferocytosis and a target for efferocytosis inhibition in this process. Efferocytosis is a potent immunosuppressive mechanism, so this process provides an additional mechanism by which large glioblastoma tumors can evade immune responses.

## INTRODUCTION

Glioblastoma is an aggressive form of brain cancer that is currently incurable. Common histological features of glioblastoma are diffuse infiltration, hypercellularity, tumour cell proliferation, nuclear pleomorphism, microvascular proliferation and geographic necrosis. The last of these, necrosis, is due to thrombotic events in the tumour microvasculature that occur as a result of the secretion of pro-coagulants by tumour cells(1). This results in the death of cells in the vicinity of the compromised vasculature along with the movement of cancer cells away from the necrotic region, which results in a “palisading” ring of cells around the region of necrosis. These necrotic regions cause a substantial remodelling of the tumour microenvironment and current evidence indicates that they promote aggressive cancer growth(2).

Glioblastoma elicits a multi-layered immunosuppression that acts both locally and systemically(3). Glioblastoma patients show marked reductions in circulating CD4+ T cells and increases in circulating myeloid-derived suppressor cells(4, 5). Local immunosuppressive effects are mediated by the glioblastoma cells themselves, which can secrete immunosuppressive cytokines such as TGFβ(6), as well as indirectly via glioblastoma recruitment of regulatory T cells and various myeloid cell types, including microglia, macrophages and myeloid-derived suppressor cells(3, 7). While engagement of myeloid cells is common in many cancers, this feature is particularly pronounced in glioblastoma, where myeloid cells sometimes comprise over 50 % of cells in tumours(8, 9). Based on microarray expression and RNA-seq studies, glioblastoma has been grouped into molecular subtypes (most commonly proneural, mesenchymal and classical) that are associated with different mutational profiles(10, 11). While all glioblastoma subtypes have a poor prognosis, mesenchymal subtype patients have a shorter median overall survival compared to other subtypes(11). The mesenchymal subtype has the highest propensity for myeloid cell recruitment, with a higher proportion of these being macrophages(11, 12). Microglia and macrophages in the glioblastoma tumour microenvironment adopt an immunosuppressive phenotype and are a major obstacle to effective immunotherapy for glioblastoma(13).

Immune function of microglia and macrophages is determined in part by their cytokine secretion profile (14) and in part by their phagocytic activity. Phagocytosis can be either immunogenic or tolerogenic. Dendritic cells are very effective at directing phagocytosis towards immunogenic antigen presentation but are rare in both normal brain and in glioblastoma tumours(15). Macrophages and microglia are also capable of antigen presentation after phagocytosis. However, compared to dendritic cells, macrophages have a reduced antigen-presenting ability due both to their inability to traffic to draining lymph nodes and their lower cross-presentation capacity (reviewed in (16)). Macrophages also engage in efferocytosis, a term that refers specifically to the phagocytosis of apoptotic cells(17). Immunologically, efferocytosis is tolerogenic. Efferocytosis sequesters apoptotic cells from other immune cells, bypasses class II MHC processing and degrades proteins more rapidly than phagocytosis, preventing antigen presentation(16). In addition, efferocytosis induces a switch in macrophage cytokine expression from pro-inflammatory cytokines to immunosuppressive cytokines(18, 19). Therefore, in the tumour microenvironment, ongoing efferocytosis potentially contributes to the establishment and maintenance of immunosuppression. Consistent with this, selective inhibition of efferocytosis in an immunocompetent colon adenocarcinoma mouse model suppressed cancer growth and also enhanced the anticancer activity of immune checkpoint inhibitors targeting PD-1 or PDL1(20).

Transglutaminases are a family of nine proteins, best known for their role in the formation of cross-linked protein polymers (21). The most widely studied of these is TGM2 (tissue transglutaminase, also referred to as TG2 or TG). TGM2 has the transglutaminase activity that defines this enzyme family, along with an atypical G protein activity(22). It is expressed both intracellularly and extracellularly; extracellular expression occurs via a cell surface translocation process, rather than by a classical leader sequence-driven export mechanism(21). TGM2 knockout mice are viable but are prone to developing autoantibodies and immune complex glomerulonephritis(23). This is due to impaired efferocytosis in TGM2-null macrophages, a defect that was evident in multiple tissues. Macrophages from TGM2-null mice bind apoptotic cells but are defective in the engulfment step, which requires a functional TGM2 GTP binding site(24). This efferocytosis defect prevents the induction of immunosuppressive cytokines in macrophages, as an inflammatory response is seen in the knockout mice, but not the wild-type mice, after induction of apoptosis(23). Induction of apoptosis in mouse thymus or liver led to an increase in TGM2 expression that was TGFβ-dependent(23). The process of efferocytosis therefore both requires TGM2 and increases TGM2 expression in macrophages.

TGM2 has been implicated in numerous diseases(25), including cancer(26), and small molecule inhibitors of its transglutaminase and GTP binding activities are under development(27). Specifically in glioblastoma, TGM2 has been found to be preferentially expressed in mesenchymal subtype glioblastoma and its inhibition in a subset of glioblastoma cells reduces proliferation in cell culture and in a xenograft model(28, 29). The role of TGM2 in glioblastoma-associated immune cells has not been studied to date. Here we show that glioblastoma-associated macrophages are a major source of TGM2 in glioblastoma. Immunohistochemical analyses in both mouse xenograft models and human samples show particularly high expression in macrophages surrounding regions of necrosis, where the macrophages are engaged in clearing away apoptotic cells by efferocytosis. When grown in physiologic oxygen concentrations, patient glioblastoma organoids actively engaged in efferocytosis and this could be blocked by TGM2 inhibition. Immune checkpoint inhibitors, while effective in other cancer types, have shown limited activity in glioblastoma (30, 31). Given the well-established immunosuppressive role for efferocytosis, blocking intratumoural efferocytosis via TGM2 inhibition may be a promising new way to sensitize glioblastoma to these immunotherapies.

## MATERIALS AND METHODS

### Bioinformatics

TCGA data were analyzed using cBioportal (32, 33) using the Cell 2013 RNAseq dataset(34). Chinese Glioma Genome Atlas (CGGA) data (35) were analyzed using GlioVis(36). Data from TCGA were also downloaded from cBioportal for analysis using Enrichr(37, 38). Single cell RNAseq data from Abdelfattah *et al.* (8) were analyzed using the Broad Institute Single Cell Portal. (https://singlecell.broadinstitute.org/single_cell).

### Antibodies

TGM2 (D11A6) rabbit monoclonal antibody (cat. # 3557), Iba1 (E4O4W) rabbit monoclonal antibody (cat. # 17198, used for immunohistochemistry), rabbit DA1E monoclonal antibody IgG isotype control (cat. # 3900) and cleaved caspase-3 rabbit monoclonal antibody (cat.# 9664) were all from Cell Signaling Technology. Anti-Iba1/AIF1 mouse monoclonal antibody (used for immunofluorescence in xenografts) and anti-CD68 mouse monoclonal antibody (used for immunofluorescence in human tissue samples) were from Millipore/Sigma (cat. #s MABN92 and AMAB90873, respectively). The anti-Iba1 rabbit monoclonal antibody used for immunofluorescence in organoids was from Cell Signaling Technology (cat. #17198). GAPDH mouse monoclonal antibody (cat. # ab8245) was from Abcam.

### Cell culture

PriGO17A cells were isolated from a glioblastoma patient undergoing a first surgery as described previously (39) following the protocol described by Pollard *et al*. (40). Glioblastoma cells were isolated and grown on laminin-coated plates in neurobasal A medium supplemented with B27, N2, EGF and FGF in 5% O_2_ and 5% CO_2_. THP-1 cells and Jurkat cells (both from ATCC) were grown in RPMI-1640 medium supplemented with 100 units/mL penicillin, 100 μg/mL streptomycin and 10% fetal bovine serum at 37°C and 5% CO_2_. To differentiate THP-1 cells into macrophages, they were treated with 20ng/mL phorbol 12-myristate 13-acetate (PMA) for 5 days(41). PMA media was then removed and replaced with regular media. Cells were routinely tested and shown to be free of mycoplasma using PCR analysis.

### CCL2 ELISA

Levels of secreted CCL2 produced by glioblastoma cells were measured by ELISA using a RayBio Human MCP-1 ELISA kit (cat. # ELH-MCP1).

### Xenograft model

Female CD1-nude mice were injected intracerebrally with PriGO17A cells using a stereotaxic apparatus as described previously(39). Mice were euthanized at the first signs of morbidity. Brains were isolated, fixed with formalin and embedded in paraffin.

### Western blotting

Western blotting was done as described previously(42). Blots were stained with amido black as a loading control prior to probing with antibody. Blots were also probed with antibody to GAPDH as an additional loading control.

### Immunohistochemistry

Immunohistochemistry was performed on formalin-fixed, paraffin-embedded tissue sections using the Leica Bond system (Leica Biosystems, Inc.). For rabbit antibodies, a modification of the Leica Bond system IHC protocol F that eliminates the post primary step when using rabbit antibodies on mouse tissue was used. Sections were first treated using an EDTA buffer (pH 9.0, epitope retrieval solution 2) for 20 min. Sections were then incubated with a 1:100 dilution of primary antibody for 30 min and detected using an HRP-conjugated compact polymer system. For the rabbit IgG control, a concentration matching the protein concentration of the specific antibody was used. Slides were then stained using DAB as the chromogen, counterstained with hematoxylin, mounted and coverslipped. Whole section digital images were generated using a Zeiss Axioscan.Z1 slide scanner.

### Immunofluorescence microscopy on tissues

For mouse xenograft samples, paraffin sections were first deparaffinized and treated using an EDTA buffer pH 9.0 for antigen retrieval. Sections were blocked for 30 minutes with Rodent Block M (Biocare, cat. # RBM961H). Sections were then incubated overnight at 4°C with either no primary antibody, 1:75 dilution of rabbit TGM2/1:500 mouse Iba1 or 1:75 dilution of rabbit TGM2/1:1000 mouse Iba1. The following day, sections were washed with 1 X TBST and incubated with goat anti-rabbit IgG-488 and donkey anti-mouse IgG-568 using a 1:500 dilution for 1h at room temperature. This was followed by incubation for 5 minutes with a quencher (Vector TrueView Autofluorescence Quenching Kit #SP-8400, Vector Labs) to decrease autofluorescence. Sections were then washed, incubated with 5 µg/ml of DAPI and coverslipped. For immunofluorescence on human glioblastoma sections, sections prepared as above were incubated with a 1:75 dilution of TGM2 antibody and a 1:1000 dilution of CD68 antibody, followed by incubation donkey anti-rabbit IgG-488 and goat anti-mouse IgG-594, both at a 1:500 dilution. Microscopy was performed using a Zeiss Axioskop 2 fluorescence microscope or a Zeiss Axioimager.M2 with Z stack acquisition.

### Tissue microarray and whole slide human glioblastoma samples

Construction of the tissue microarray was described previously (43). Duplicate 1 mm cores were used for each patient. While the original tissue microarray included lower grade glioma patients, only tissue from the eighty-three IDH wild-type glioblastoma patients was analyzed here. Whole sections from four patients were also assessed as the tissue microarray cores generally lacked necrotic regions. Immunohistochemistry and immunofluorescence were performed as described above.

### Mouse macrophage isolation

Monocytes were isolated from the bone marrow of C57BL/6 mice and differentiated into macrophages by incubation for seven days in media containing recombinant CSF1 (R&D Systems).

### Immunofluorescence microscopy for TGM2 during efferocytosis in cell culture

THP-1 cells were grown on coverslips in 6 well plates and differentiated with PMA as above. PMA was removed after five days and replaced with fresh media. Jurkat cells were treated with 1 µM staurosporine for 3 h to induce apoptosis (confirmed by microscopy with fluorescent Annexin V), washed, resuspended in media and added to the THP-1 cells. At the indicated timepoints, media was removed and cells were fixed with paraformaldehyde. Immunofluorescence was performed using TGM2 antibody at a 1:200 dilution. Quantitation of TGM2 immunofluorescence was done using Fiji software. TGM2 expression in bone marrow-derived macrophages during efferocytosis was determined using the same procedure.

### Efferocytosis assay

Efferocytosis was assayed following the protocol described above for TGM2 immunofluorescence described, except that after staurosporine treatment Jurkat cells were labelled with pHrodo following the kit manufacturer’s instructions. In addition, THP-1 cells were pretreated with either ONO-7475 (final concentration 5 nM, added at least 45 min before the addition of apoptotic cells) or NC9 (final concentration 10 µM, added 24 h before the addition of apoptotic cells) in media. At indicated time points, media was removed and cells were fixed with paraformaldehyde. pHrodo fluorescence was assessed by fluorescence microscopy and analysis of digitized images using Fiji software (44).

### Organoid culture and analysis

Glioblastoma organoids were generated and cultured as described by Jacob *et al*. (45), except that cultures were maintained in 5% O_2_ unless otherwise indicated. Organoids were analyzed between two and four weeks after isolation from patients and were treated with either 5 nM ONO-7475 or 10 µM NC9 for one or two days before fixation with 4% paraformaldehyde. TUNEL assays were performed using the Click-iT™ Plus TUNEL Assay Kits for In Situ Apoptosis Detection (ThermoFisher cat. # C10617) following the manufacturer’s recommendations. Immunofluorescence for Iba1 (Cell Signaling Technology 1:200) was performed after this. Microscopy and image analysis were as above.

### Statistics

Graphing and statistical analyses were done using SigmaPlot 14.5. Details of the statistical tests used are given in the Figure legends.

## RESULTS

### Expression in glioblastoma subtypes

Analysis of TGM2 mRNA levels in the TCGA 2013 glioblastoma dataset (152 patients) using cBioportal showed significantly higher expression in the mesenchymal subtype (Figure 1A). This was also observed in the Chinese Glioma Genome Atlas (CGGA; 249 patients) database analysed using GlioVis (Figure 1B). As the mesenchymal subtype is known to have higher infiltration of myeloid cells (microglia, macrophages and/or myeloid-derived suppressor cells) relative to other glioblastoma molecular subtypes, correlations between mRNA levels of TGM2 and CD68, a standard microglia/macrophage marker, were assessed. TGM2 mRNA had a positive linear correlation with CD68 mRNA in both the TCGA and CGGC datasets (Figure 1C and D). As a second way to assess whether high TGM2 was associated with high levels of macrophage/microglia, a TGM2 gene expression signature was generated by identifying 50 genes with expression levels that had the highest positive correlation with TGM2 mRNA levels in the TCGA dataset. This expression signature was then analyzed in Enrichr under “cell types”. This gave a very strong match with macrophages (Figure 1E; 34/50 match between TGM2 signature and macrophages; adjusted *P* = 2.8×10^−15^, odds ratio 5.35). This suggests that macrophages are a major source of TGM2 mRNA in glioblastoma. TGM2 mRNA expression was also assessed in the single cell RNA-seq dataset from Abdelfattah *et al*. (8) that is available through the Broad Institute Single Cell Portal. Figure 1F shows Uniform Manifold Approximation and Projection (UMAP) clustering with cell type assignments on the left and TGM2 expression mapped onto these in the adjacent plot. This showed that highest TGM2 mRNA expression was in myeloid cells and endothelial cells (Figure 1F). Figure 1G shows the distribution of TGM2 expression in the assigned clusters described in Abdelfattah *et al*.; endothelial cells consistently have high TGM2 expression, while only a subset of myeloid have high expression. With the immune cell subclustering used in Abdelfattah *et al*., TGM2 expression was highest in the s-mac 2 cluster, corresponding to macrophages expressing immune-suppression markers (Figure 1H). High expression was also observed in myeloid-derived suppressor cells and M2-like macrophages (s-mac-1 cluster expressing M2-like macrophage markers). Directly relevant to further studies below, these subclusters all express a hypoxia signature(8). TGM2 mRNA expression was also present in a subset of glioblastoma cells, in agreement with previous literature, although this is lower than the expression in either endothelial cells or myeloid cells. Overall, these data indicate that macrophages are a major source of TGM2 mRNA in glioblastoma tumours.

**Figure 1.**
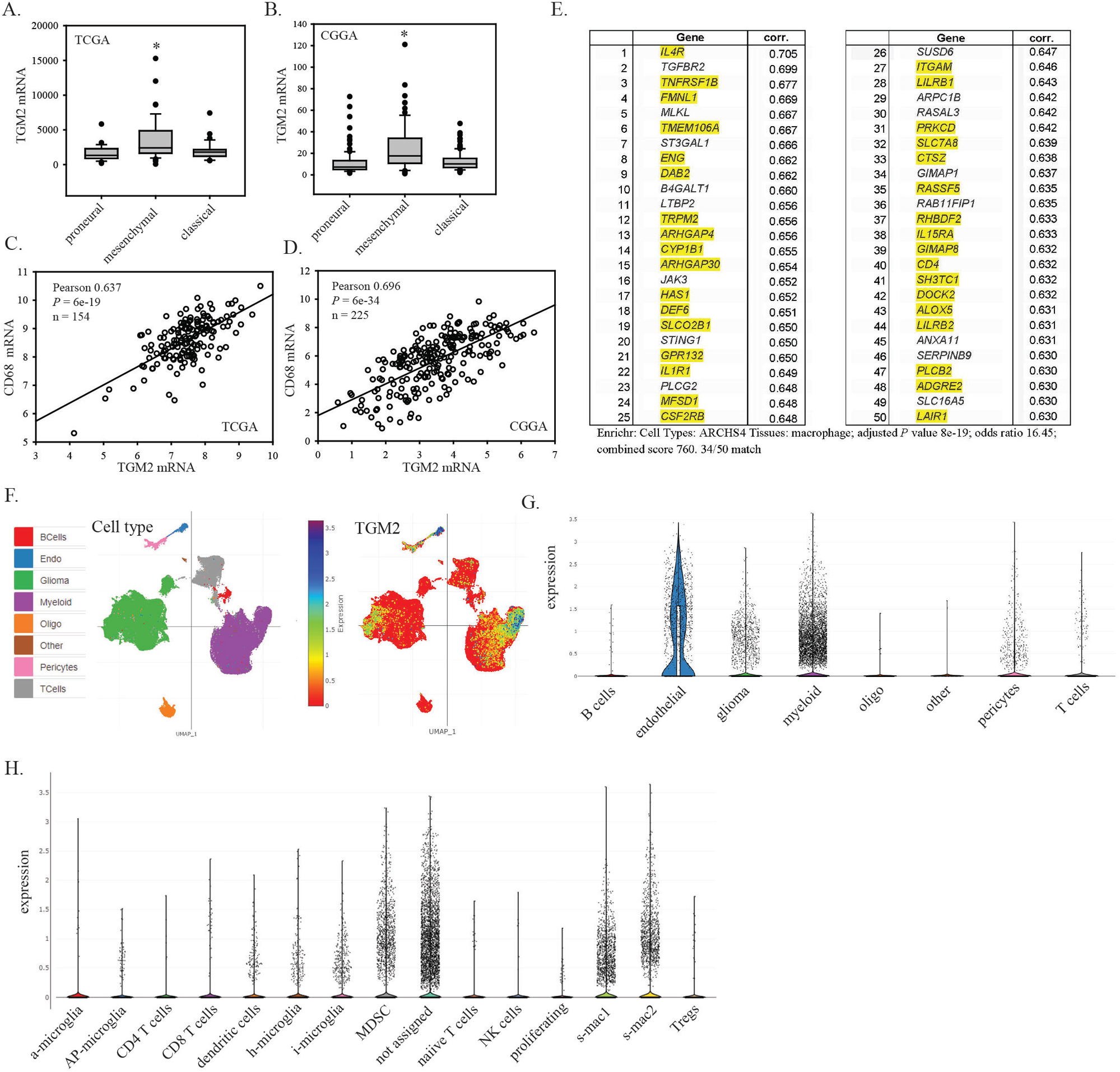
TGM2 mRNA expression in glioblastoma. A and B. TGM2 RNA expression in glioblastoma molecular subtypes in the TCGA and CGGA datasets; C and D. Correlation of TGM2 mRNA levels with mRNA levels for the macrophage/microglia marker CD68 in the TCGA and CGGA datasets; E. The fifty genes showing the highest positive Spearman correlation with TGM2 mRNA expression in the TCGA database were identified using cBioPortal. This gene set was then analyzed for matches to cell type datasets. Yellow highlights show the match to macrophage cell type identified using Enrichr under Cell Types/ARCHS4 Tissues (34/50 match, qval = 2.8×10-15, odds ratio 5.35); F. TGM2 mRNA expression in single cell RNAseq data. Plots show UMAP clustering using data from Abdelfattah *et al*. analyzed with the Broad Institute Single Cell Portal. Plots show cell type assignments (left) and TGM2 mRNA expression (right); G. Violin plots (with all data points) of TGM2 mRNA expression in glioblastoma tumour cell types; H. Violin plots (with all data points) of TGM2 mRNA in the glioblastoma tumour immune cell subtypes described by Abdelfattah *et al.*

### Mesenchymal-subtype glioblastoma xenograft model

To model mesenchymal-subtype glioblastoma in a xenograft model, we used PriGO17A glioblastoma cells that we isolated previously from a patient undergoing a first surgery for glioblastoma. These cells were isolated using serum-free conditions and growth in 5% O_2_ on laminin-coated plates, as described previously(39, 40). Microarray expression analysis showed that these were mixed mesenchymal/classical subtype(46). After intracerebral injection, these cells formed tumours in immunocompromised mice that showed marked engagement and activation of mouse macrophage/microglia in regions adjacent to and within the tumour microenvironment (Figure S1A-D). PriGO17A cells have increased expression of CCL2 mRNA (Figure S1E) and secrete high levels of CCL2 protein (Figure S1F) relative to three predominantly classical subtype glioblastoma cell cultures isolated from different patients under identical conditions. Human CCL2 is a well-known macrophage chemoattractant that is expressed at higher levels in mesenchymal subtype glioblastoma(47). As human CCL2 is also active in mice, this provides a likely explanation for the strong recruitment of macrophages in this model.

### Immunohistochemical analysis of TGM2 expression in normal and xenografted mouse brain

A rabbit monoclonal antibody to TGM2 was chosen to avoid mouse-on-mouse artefacts in immunohistochemistry and to ensure reproducibility. This antibody labels predominantly a single band of the expected size on Western blots of macrophage extracts; this band is absent in PriGO17A glioblastoma cells (Figure S2A-C). Immunohistochemistry for TGM2 in normal mouse brain (Figure 2A) showed positive staining that was absent in no primary controls and non-specific primary rabbit antibody controls (Figure S2D). Microglia were weakly positive (Figure 2B). Endothelial cells in normal brain vasculature were strongly positive (Figure 2C). Positive TGM2 staining was also seen in the pia mater and ependymal cells lining the lateral ventricle (Figure 2D and E).

**Figure 2.**
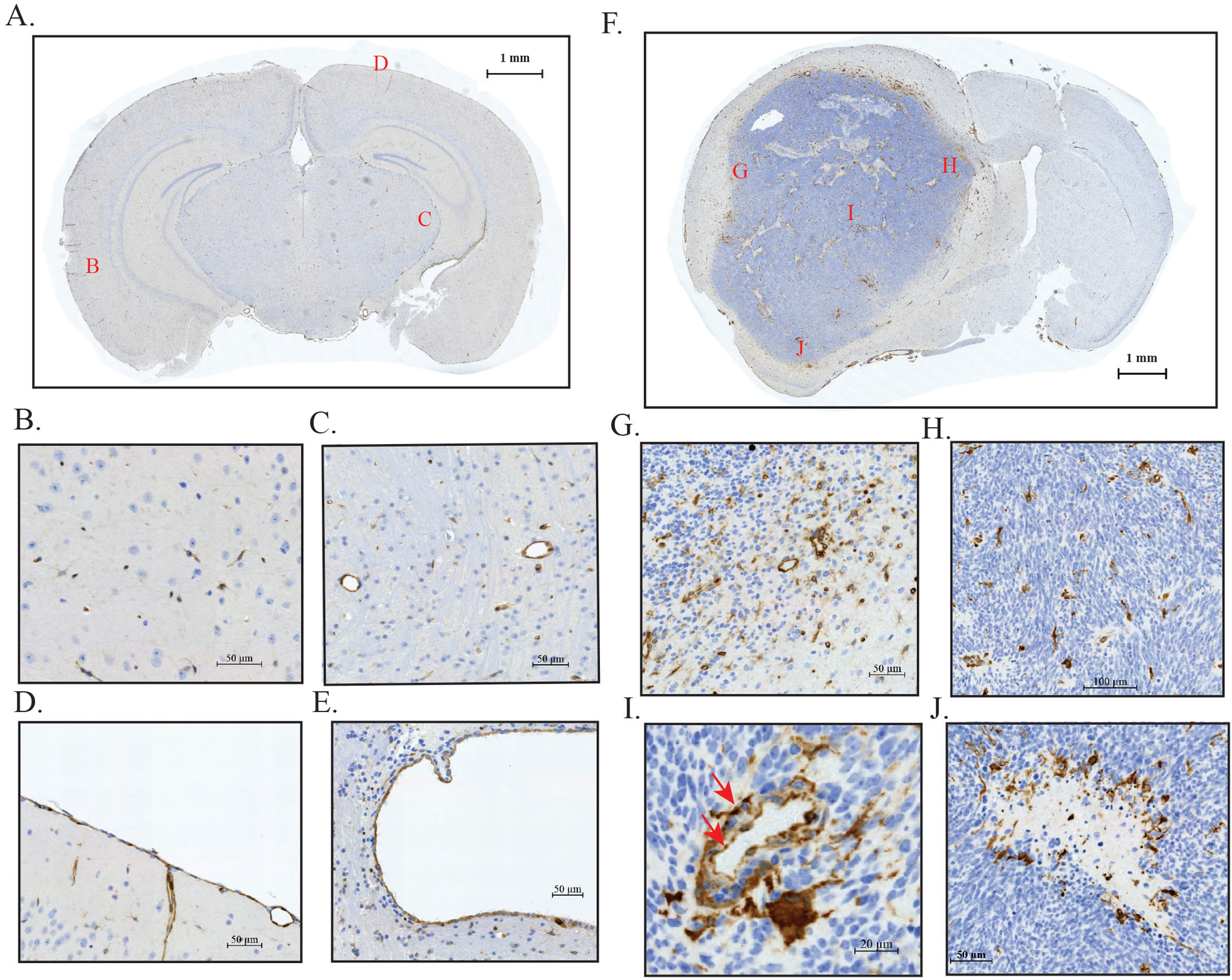
TGM2 expression in normal and human glioblastoma xenograft mouse brain. A. Image of normal whole mouse brain section stained for TGM2. Red letters indicate regions for enlarged areas shown in B-E; B. TGM2 staining in brain parenchyma; C. TGM2 staining in brain endothelial cells; D. TGM2 staining in pia mater; E. TGM2 staining in ependymal cells (from section shown in F, non-injected hemisphere); F. PriGO17A cells were injected intracerebrally into CD1 nude mice. Mice were euthanized when they showed signs of morbidity and brains were formalin-fixed and paraffin-embedded. An image of a whole brain section stained with TGM2 antibody is shown. Red letters indicate regions for the enlarged images shown in G-J. G. Diffusely infiltrating tumour margin; H. TGM2 staining of microglia/macrophages in central tumour mass; I. TGM2 staining of tumour endothelial cells (red arrows); J. TGM2 staining of microglia/macrophages bordering and within a necrotic region.

Figure 2F shows a coronal brain section of an immunocompromised mouse that was injected intracerebrally with PriGO17A cells. The large tumour has diffusely infiltrating margins (Figure 2G), hypercellularity (Figure 2H), abnormal vessels (Figure 2I), and geographic necrosis (Figure 2J), recapitulating common pathological features of human glioblastoma. TGM2 is not detected in PriGO17A glioblastoma cells. However infiltrating macrophages are strongly positive (Figure 2G and H). As with normal brain vasculature, endothelial cells of the tumour vasculature also express TGM2 (Figure 2I). The strongest staining is seen in macrophages surrounding and within regions of necrosis (Figure 2J). Macrophages in these images were identified by morphological criteria. To confirm their identity, double immunofluorescence for Iba1 and TGM2 was performed on sections. There was a clear overlap in staining between the two markers, confirming the identity of TGM2 positive cells as either macrophages or microglia (Figure S3).

As mentioned earlier, necrotic regions in glioblastoma arise as a consequence of thrombotic events in the abnormal vasculature. These necrotic regions display a characteristic “serpiginous” or “geographic” architecture. A likely function for macrophages in this setting is the phagocytosis of dead and dying cells. Figure 3A and B show two different regions of geographic necrosis in the xenograft model after staining for TGM2. The presence of high TGM2-expressing macrophages around the edges and within these regions of geographic necrosis is a very consistent feature. Figure 3A shows a necrotic region that has a large number of apoptotic cells within it. Palisading glioblastoma cells, interspersed with apoptotic cells, border the necrotic region. High TGM2-expressing macrophages are present within the necrotic region. Higher magnification images of these show morphologies consistent with active efferocytosis (Figure 3C). Figure 3B shows a necrotic region in which apoptotic cells are largely absent from the central region. Macrophages line the rim of the necrotic region and also appear to be engaged in efferocytosis (Figure 3D). Their large size is probably a consequence of their having engulfed multiple apoptotic cells. Apoptotic cells were identified by morphological criteria (condensed nuclei) in these images. To support this, we also performed immunohistochemistry with antibody to cleaved caspase-3. Cleaved caspase-3 positive cells were present in regions of necrosis (Figure 3E and F). Some of these were observed to be in clusters around a normal nucleus, suggestive of macrophage uptake of multiple apoptotic cells (Figure 3G).

**Figure 3.**
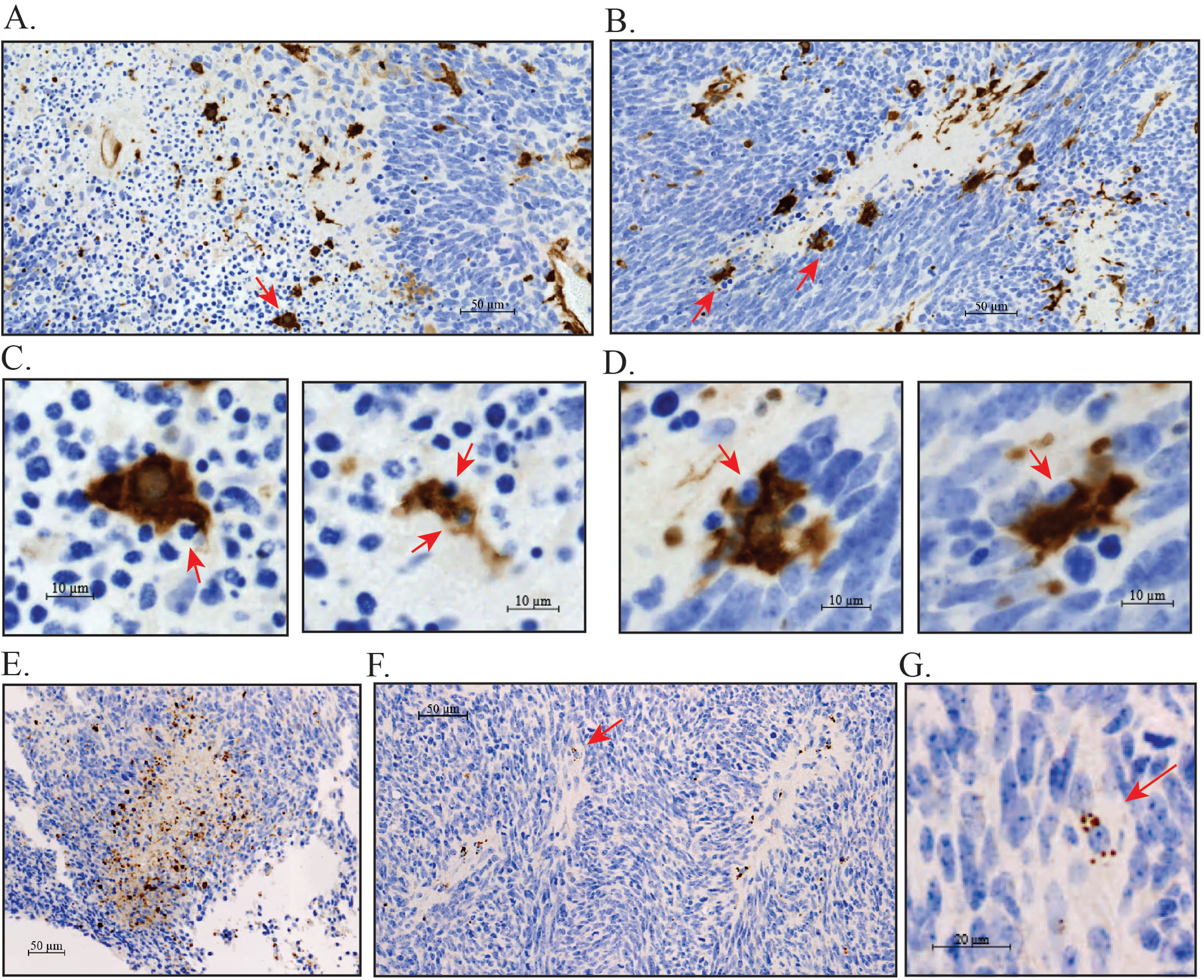
TGM2 staining in necrotic regions of the xenograft. A. Necrotic region showing palisading (region under red bracket). Abundant apoptotic cells are present within the necrotic region and between the palisading cells (small dark blue nuclei, examples indicated with red arrowheads). Strongly TGM2 positive cells are present within the necrotic region; B. Necrotic region with few apoptotic cells. Strongly TGM2-positive cells are predominantly localized around the edges of the necrotic region. For A and B, red arrows indicate TGM2-positive cells showing apparent efferocytosis of apoptotic cells; C. Examples of TGM2-positive cells engaged in efferocytosis within necrotic regions. Left panel shows closeup of cell indicated by red arrow in A, right panel shows example taken from outside the area shown in A; D. Examples of TGM2-positive cells engaged in efferocytosis at the edges of necrotic regions. Closeups are of cells indicated with red arrows in B; E. Region of necrosis in mouse xenograft model with immunohistochemistry for cleaved caspase 3. Multiple cleaved caspase 3-positive apoptotic cells are present within and around the border of the necrotic region; F. Smaller regions of necrosis showing cleaved caspase 3-positive apoptotic cells; G. Cleaved caspase 3-positive apoptotic cells clustered around a normal cell nucleus, consistent with macrophage efferocytosis (closeup of a cell marked with a red arrow in F.).

### TGM2 expression in glioblastoma patients

To determine if TGM2 showed similar expression patterns in human glioblastoma, we performed TGM2 immunohistochemistry on whole sections from patients. Figure 4A shows a low magnification image of a patient section in which three necrotic regions are visible. Multiple cells with strong TGM2 staining are present in the vicinity of the necrotic regions. Figure 4B shows a higher magnification image of a necrotic region with strongly TGM2 positive cells lining the region, very similar to what was observed in the xenograft model. To confirm that these cells are macrophages, we performed double immunofluorescence for TGM2 and the macrophage marker CD68. Imaging was performed using Z stack acquisition. As expected, endothelial cells were TGM2-positive but CD68-negative (Figure 4C-E). Outside of endothelial cells, CD68 and TGM2 immunofluorescence co-localized (Figure 4C-F). This confirms that the high TGM2 expression seen in cells lining necrotic regions is in macrophages, as observed in the mouse model. Macrophage CD68 immunofluorescence was punctate (Figure 4C-F), consistent with its previously described endosomal/lysosomal subcellular location (48). TGM2 immunofluorescence in macrophages was also punctate (Figure 4C-F). This punctate pattern is also present in the immunohistochemistry analyses but is not always as apparent because of the lower dynamic range. CD68 and TGM2 therefore also co-localize at a subcellular level, suggesting that the very highest TGM2 expression is also found in the endosomal/lysosomal compartment. TGM2 expression was also assessed by immunohistochemistry on a tissue microarray containing duplicate 1 mm cores from 84 patients. Examples of the immunohistochemistry are shown in Figure S4A-D and the scoring is in Table S1. Although the cores generally did not have significant necrosis (likely due to selection bias in choosing areas to be cored), strong positive TGM2 cells were occasionally seen in focal areas in the tissue microarray samples. Sporadic cells with strong positive TGM2 staining that appeared to be engaged in efferocytosis were also observed (Figure S4D). Tumour cells were generally negative (Table S1).

**Figure 4.**
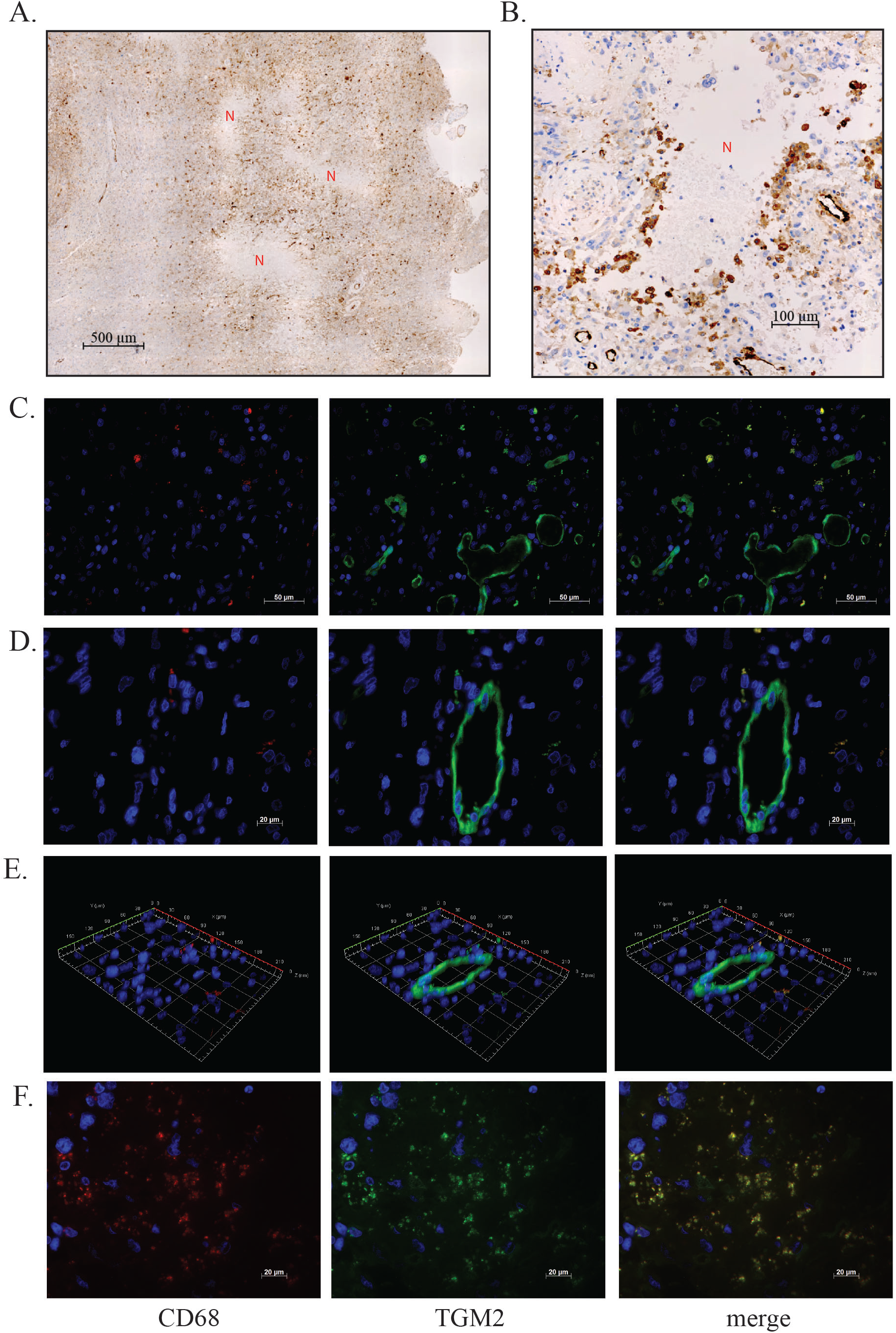
TGM2 staining patterns in human glioblastoma tumours. A. Image from full section showing strong TGM2 expression near regions of necrosis (indicated with N). Regions without necrosis (lower left) show lower levels of TGM2 staining; B. Necrotic region showing TGM2 positive cells surrounding the border of the necrotic region, similar to what was observed in the xenograft model. C-F. Double immunofluorescence for CD68 and TGM2 in whole sections from patient tumours. For each image row, the left panel shows CD68, the middle panel shows TGM2, and the right panel shows the merged image. Except for E, each row shows a single focal plane image from the Z stack. Row C shows a region with macrophages and endothelial cells. Row D shows a higher magnification view of a tumour blood vessel with adjacent macrophages to illustrate differences in subcellular location. Row E shows a Z stack of the same region as B, emphasizing the co-localization of TGM2 and CD68 in macrophages. Row F shows an area adjacent to necrotic regions.

### TGM2 expression in macrophages undergoing efferocytosis

The above findings suggest that high TGM2 expression is a marker of efferocytic macrophages. To explore this further, we assessed TGM2 expression in efferocytic macrophages in cell culture. Macrophages were generated by phorbol myristate acetate-induced differentiation of THP-1 human monocyte cells. Staurosporine-treated Jurkat T cells were used as a source of apoptotic cells and were added to the differentiated THP-1 cells to initiate efferocytosis. Exposure of macrophages to apoptotic cells caused an increase in intracellular TGM2 levels that was highest after 4 h, the longest time point assessed (Figure 5A and B). A similar result was observed when primary mouse macrophages were substituted for THP-1 cells in the same assay (Figure 5C). In this assay, internalization of apoptotic cells peaks after approximately 1 h and digestion has typically taken place by 4 h. The increase in TGM2 expression therefore occurs later in the process of efferocytosis. These cell culture findings are consistent with high TGM2 expression functioning as a marker of efferocytic macrophages.

**Figure 5.**
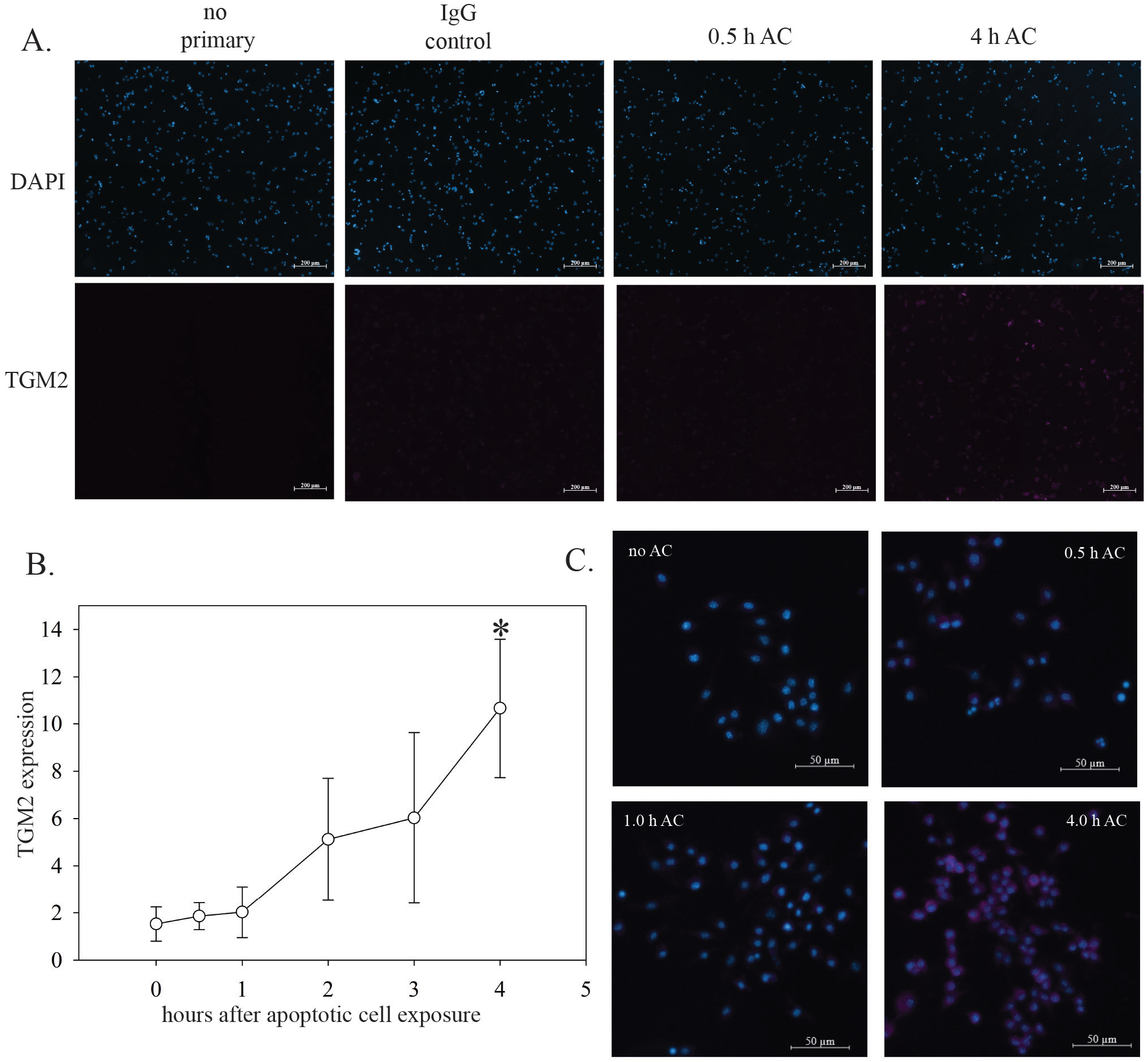
TGM2 expression during macrophage efferocytosis in cell culture. A. THP-1 cells grown on coverslips were differentiated into macrophages and then exposed to apoptotic Jurkat T cells (AC). At the indicated time, cells were fixed in paraformaldehyde and immunofluorescence for TGM2 was performed. No primary and non-specific rabbit IgG controls for immunofluorescence staining are shown. B. Quantitation of TGM2 immunofluorescence. Data are from three biological replicates. Error bars show standard deviation. * indicates P < 0.05 by a two-sided t test relative to the 0 h value. C. Mouse macrophages derived from bone marrow were exposed to apoptotic cells and immunofluorescence for TGM2 was performed as in A.

### Small molecule TGM2 inhibitors block efferocytosis

Several TGM2 inhibitors were tested to determine if TGM2 could be targeted pharmacologically for efferocytosis inhibition. As above, this was assayed using differentiated THP-1 cell macrophages and staurosporine-treated Jurkat T cells as a source of apoptotic cells. The latter were labelled with the pH-sensitive dye pHrodo to monitor apoptotic cell internalization into acidified vesicles during efferocytosis. The dual MERTK/AXL tyrosine kinase inhibitor ONO-7475 (49, 50) was used as a positive control, given the well-established role for these receptors in efferocytosis. ONO-7475 inhibited efferocytosis at a concentration of 5 nM, a concentration where it is selective for MERTK and AXL (Figure 6A). The irreversible TGM2 inhibitor NC9 (51) also inhibited efferocytosis, showing a slightly higher overall inhibition than observed with ONO-7475 (Figure 6A). This was tested at 10 µM, a lower concentration than has been used in previous studies to inhibit intracellular targets. We also tested NCEG2, a recently described irreversible TGM2 inhibitor that was specifically designed to be cell-***im***permeable(52). This also inhibited efferocytosis at a concentration of 10 µM (K_I_ for purified TGM2 enzyme is 4.1 µM) (Figure 6A).

**Figure 6.**
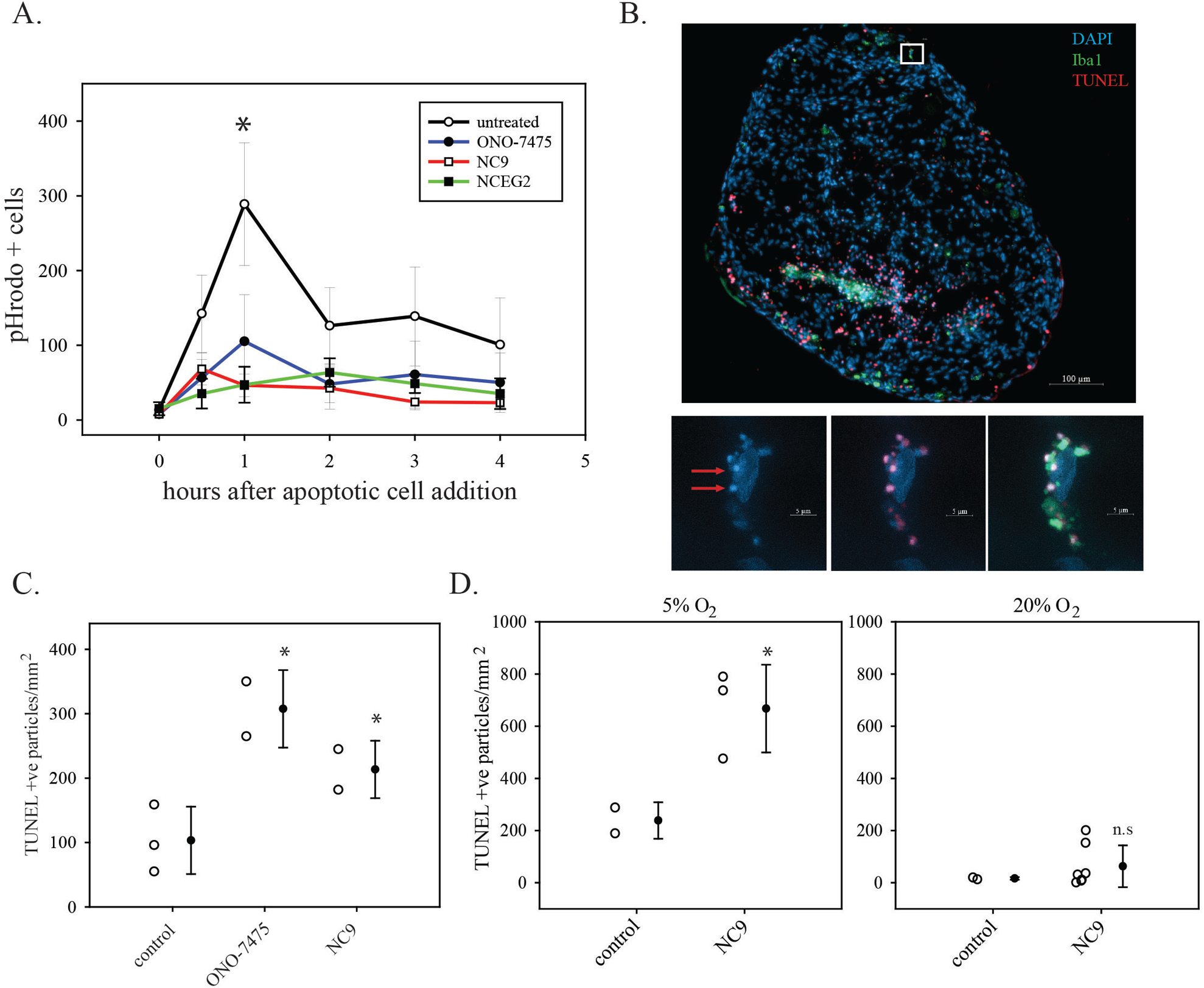
Inhibition of efferocytosis by TGM2 inhibitors in cell culture and in organoids. A. THP-1 cells growing on coverslips were differentiated into macrophages and exposed to apoptotic cells that were labelled with pHrodo. Cells were fixed at the indicated times and analyzed for pHrodo fluorescence by microscopy. Numbers of positive cells were quantitated using Fiji software. Data shown are the mean and standard deviation from three biological replicates. For each biological replicate, 3-4 random fields of view were analyzed. * indicates P < 0.05 by a two-sided t test in comparisons of the untreated 1 h time point data with either the ONO-7475 or the NC9 1 h time point data. B. Efferocytosis in human glioblastoma organoids. Nuclei are labelled with DAPI (blue); macrophages/microglia are detected with Iba1 immunofluorescence (green); apoptotic cells are detected by fluorescent TUNEL assay (red). The small white square indicates the site of the cell shown in close up in the bottom images. Bottom images show a macrophage/microglial cell efferocytosing multiple apoptotic cells. Left panel shows DAPI only; middle sample shows DAPI and TUNEL; right panel shows DAPI, TUNEL and Iba1. Red arrows indicate examples of apoptotic cells with nuclear condensation. C. Quantitation of apoptotic cells in organoids that were either untreated (control) or treated with 5 nM ONO-7475 for 48 h or 10 µM NC9 for 24 h. Apoptotic cells were counted using image analysis and were normalized to organoid surface area, as described in Materials and Methods. Circles show apoptotic cells counts from two or three sections from one organoid per condition. Mean and standard deviation are shown in the adjacent closed circle with error bars. * indicates a P value < 0.05 using a one-tailed Student’s t-test. n.s. not significant. D. Organoids from a second patient were isolated using either 5% O2 or 20% O2 conditions. Once established, organoids were either untreated or treated with 10 µM NC9 for 48 h. Quantitation of apoptotic cells was performed as in C. Each open circle shows the apoptotic cell count from one organoid. Mean and standard deviation are shown in the adjacent closed circle with error bars. * indicates a P value < 0.05 using a two-tailed Student’s t-test. n.s. not significant

### Inhibition of TGM2 in patient-derived organoids

Recently, methods for the long-term culture of patient glioblastoma samples as organoids have been developed (45). Along with the glioblastoma cells, these organoids maintain native populations of immune cells including myeloid cells, making them a potentially valuable tool for studying processes such as efferocytosis in an environment that closely models the human glioblastoma tumour microenvironment. TUNEL assays for apoptotic cells, along with immunofluorescence for Iba1 (to label macrophages) and DAPI counterstaining were used to assess efferocytosis in glioblastoma organoid samples. Organoids showing high cellularity were used for analyses (the culturing process also yields tissue pieces that are primarily matrix with few or no viable cells). Figure 6B shows an untreated patient organoid patient after TUNEL labelling and Iba1 immunofluorescence with DAPI counterstaining. Iba1-positive cells were detected; this was observed in all organoids examined. TUNEL positive cells were also observed consistently; very bright TUNEL positive cells were frequently seen in the margin of the organoids and internal clusters of TUNEL-positive cells were seen sporadically (Figure S4E). Iba1-positive cells associated with most apoptotic cells and were observed to be engulfing apoptotic cells, which were identified both by nuclear condensation and TUNEL positivity (Figure 6B). In some, clustering of Iba1 at phagocytic portals was clearly seen, as described previously(53). Inhibition of efferocytosis is expected to lead to an increase in apoptotic cells(20). Organoids from the same patient, grown in culture for the same length of time, were either left untreated or treated with either 5 nM ONO-7475 for 48 h or 10 µM NC9 for 24 h. Organoids were then fixed and analyzed using TUNEL assays and Iba1 immunofluorescence. Both ONO-7475 and NC9-treated appeared to have reduced association of Iba1-positive cells with apoptotic cells (Figure S4E). Apoptotic cells were counted using image analysis software and counts were normalized to organoid section surface area. Treatment with both ONO-7475 or NC9 resulted in an increase in apoptotic cells (Figure 6C). To rule out possible sampling effects and to determine if the same effects were observed in other patients, organoids from a second patient were tested. NC9 treatment for 48 h also caused an increase in apoptotic cell counts in these organoids (Figure 6D). Detection of efferocytosis in these organoids requires that there is a baseline generation of apoptotic cells. The same assay was repeated using organoids that were derived at the same time from the same patient but cultured in 20% O_2_. Baseline levels of apoptosis were much lower under this condition and, consistent with the lack of apoptotic cells, efferocytosis inhibition had no effect (Figure 6D).

## DISCUSSION AND CONCLUSIONS

Analysis of multiple databases shows that myeloid cells are a major source of TGM2 mRNA in glioblastoma. Within the myeloid category, very high TGM2 mRNA expression is present in a subset of myeloid-derived suppressor cells and macrophages. High TGM2 mRNA was also present in tumour endothelial cells. This same pattern of high TGM2 expression in myeloid cells and endothelial cells was also observed in immunohistochemical analyses of a patient xenograft model and in human glioblastoma samples. These analyses also provided spatial information for the high TGM2-expressing myeloid cells, showing that they were most frequently found in the vicinity of necrotic regions. The single cell RNA-seq data from Abdelfattah *et al*. showed that the three myeloid cell subtypes that express high TGM2 mRNA (MDSC, Smac-1 and Smac-2) all have a hypoxia gene expression signature. This may be due to their proximity to necrotic regions, which are known to be hypoxic (54). TGM2 levels may be underrepresented in current datasets, as necrotic regions are often avoided during sample selection.

In the xenograft model, apoptotic cells were detected in necrotic regions using the criteria of nuclear condensation and cleaved-caspase 3 positivity. The numbers of apoptotic cells may be abnormally high in the xenograft model as the nude mice used lack Tregs, which enhance apoptotic cell clearance by macrophages(55). Apoptotic cells have previously been identified in human glioblastoma necrotic regions using both TUNEL assays and cleaved-caspase 3 immunohistochemistry(54, 56). In the xenograft model, high TGM2-expressing macrophages appeared to be engaged in clearance of apoptotic cells, as judged by both their association with areas of high apoptotic cell concentrations (potentially a consequence of the chemotactic stage of efferocytosis) and microscopic evidence for engulfment. Similar microscopic evidence for engulfment was also observed in human samples. In cell culture, exposure of macrophages to apoptotic cells induced a rise in TGM2 protein levels, supporting the concept that high TGM2 levels are indicative of efferocytic macrophages. The rise in TGM2 occurred relatively late in the efferocytosis process (maximal at 4 h after exposure to apoptotic cells), suggesting it may be an aspect of the regeneration and upregulation of efferocytosis regulators that is observed in macrophages that have already completed an initial round of efferocytosis (57).

As mentioned in the Introduction, previous work using TGM2 knockout mice showed that TGM2 has a functional role in efferocytosis(48). Specifically, cell surface TGM2 was found to promote the formation of the phagocytic portal during efferocytosis initiation(24). As a first step to determine if TGM2 has a functional role in glioblastoma efferocytosis, the ability of TGM2-selective inhibitors to inhibit efferocytosis was tested in cell culture. TAM receptor tyrosine kinases (MERTK, AXL and TYRO3) are key mediators of efferocytosis, recognizing cell surface phosphatidylserine on apoptotic cells via bridging proteins and promoting engulfment(58). A TAM receptor inhibitor was also tested in parallel with the TGM2 inhibitors. The two TAM receptors that are predominant in glioblastoma-associated macrophages are MERTK and AXL(59). The TAM inhibitor ONO-7475 (tamnorzatinib) was chosen as it has very potent activity against both MERTK and AXL (IC_50_ of 1.0 nM and 0.7 nM, respectively). ONO-7475 inhibited efferocytosis in cell culture at a low concentration (5 nM) that ensured its selectivity for MERTK and AXL. The irreversible TGM2 inhibitor NC9 also blocked efferocytosis in cell culture. A second TGM2 inhibitor, designed to be cell-impermeable (52), also inhibited efferocytosis; this is consistent with the previous work showing that cell-surface TGM2 is required to initiate efferocytosis(24). With this validation of ONO-7475 and NC9 as effective inhibitors of efferocytosis in cell culture, the next step was to evaluate their activity in a more clinically relevant context.

An ongoing challenge in testing therapeutic strategies directed against glioblastoma-associated myeloid cells is accurately modelling these cells in systems that are amenable to drug testing. To address this issue, patient-derived organoids, isolated and grown as described by Jacob *et al*.(45) were used here. When used relatively early after isolation, these maintain endogenous myeloid cell populations along with a growing population of glioblastoma cells. Analysis of these organoids with Iba1 immunofluorescence confirmed the presence of microglia/macrophages. Iba1 immunofluorescence combined with apoptotic cell labelling with a fluorescent Click chemistry TUNEL assay demonstrated that there were apoptotic cells present in the organoids and that these were being efferocytosed by the endogenous microglia/macrophages. NC9 treatment of organoids from two different patients resulted in an increase in apoptotic cells, showing that TGM2 inhibition in this clinically relevant context inhibited efferocytic clearance of apoptotic cells.

The above studies were done in organoids that were grown in 5% O_2_; this was chosen as it more closely resembles O_2_ levels in the brain(60) than 20% atmospheric oxygen. In parallel with the experiments done on the second patient organoids in 5% oxygen, effects of 20% O_2_ on apoptosis and efferocytosis were also assessed. Under these conditions, very little apoptosis was observed and, as expected in the absence of apoptotic cell generation, inhibition of efferocytosis had little or no effect. In 5% O_2_, apoptotic cells were primarily present near the margin of organoids, rather than in the central region of the organoid which is expected to have the least accessibility to O_2_. It was previously demonstrated that in organoids cultured by this method, cell proliferation primarily takes place in the margin(45). One possibility is that dividing cells may be more susceptible to apoptosis under conditions of oxygen deprivation, due to their switch to aerobic glycolysis and the accompanying increase in oxidative phosphorylation (61, 62)).

An overall model for explaining these results begins with induction of thrombosis in the tumour vasculature initiated by tissue factor secreted by the tumour cells. The subsequent hypoxia promotes death of cancer cells, which are then efferocytosed by macrophages that were previously perivascular. This efferocytosis both requires TGM2 and induces TGM2 expression. Macrophages that perform efferocytosis are known to adopt an immunosuppressive phenotype, secreting TGFβ and IL10. This change is in part mediated by epigenetic changes(63). Efferocytic macrophages also are induced to proliferate, which is proposed to amplify their immunosuppressive activity further(64). Thus, glioblastoma cells, once a tumour reaches a larger size, activate this powerful immunosuppressive mechanism to enhance their ability to avoid systemic immune responses. A previous small trial showed a small survival benefit in glioblastoma patients given immune checkpoint inhibition prior to a second surgery when they still had significant tumour burden(31). The induction of intratumoral efferocytosis may be a key factor limiting patient responses in this setting.

The data here show that efferocytosis is active in glioblastoma and can be suppressed in clinically-relevant organoid models by TGM2 inhibition. A limitation here is that current TGM2 inhibitors do not cross the blood-brain barrier. Options are either to design inhibitors that have this capacity, employ strategies to transiently open the blood-brain barrier (*e.g.* ultrasound-based methods) or use delivery methods that bypass the blood-brain barrier (*e.g.* Ommaya reservoir or other surgically-implanted slow release methods). The expression of TGM2 in normal vasculature, ependymal cells and the pia mater may also raise safety concerns with respect to TGM2 inhibition; however, the TGM2 knockout mouse does not show any obvious vascular or brain defects, perhaps because TGM2 has redundant activities with other transglutaminase family members in these tissues. Future experiments should assess TGM2 inhibitor delivery strategies in clinically-relevant animal models of glioblastoma, along with efficacy experiments as a single agent and in combination with standard therapy and immune checkpoint inhibition.

## Supporting information

Lui et al 2024 Supplemental

## ACKNOWLEDGEMENTS

We gratefully acknowledge Histology/Imaging/Staining services provided by the Louise Pelletier Core Facility (RRID: SCR_021737) at the Department of Pathology and Laboratory Services, Faculty of Medicine, University of Ottawa.

## CONFLICT OF INTEREST

The authors declare that they have no conflict of interest.

## AUTHOR CONTRIBUTIONS

ML performed the organoid culture and the immunofluorescence microscopy analyses of these, assisted with cell culture efferocytosis assays, and assisted with data acquisition and analysis for additional experiments. FS performed the cell culture efferocytosis assays. ME contributed to the design of the studies and performed the cell culture and Western blot analyses. DGM oversaw the construction and data collection for the tissue microarray used in the study. JWK initiated the study and provided expertise in TGM2 that was used in the design of experiments and the preparation of the manuscript. JS and DC contributed to the organoid studies. JW oversaw the interpretation of immunohistochemical and immunofluorescence analyses and contributed to the writing of the manuscript. IAJL oversaw the design and execution of the study, data interpretation and the writing of the manuscript. All authors reviewed the manuscript and provided comments during the writing process.

## ETHICS APPROVAL AND CONSENT TO PARTICIPATE

Isolation and use of patient glioblastoma cells and organoids was performed under Ottawa Health Sciences Network Research Ethics Board approved protocol 20120166-01H. Patient glioblastoma tissue studies were performed in accordance with the ethical guidelines of the Ottawa Hospital Research Institute and St. Michael’s Hospital.

## FUNDING

This work was funded by a project grant from the Canadian Institutes of Health Research (#162180) awarded to JWK (principal applicant) and IAJL (co-applicant).

## DATA AVAILABILITY STATEMENT

Datasets analyzed during the current study are available in from the following sources: cBioportal https://www.cbioportal.org/; Gliovis http://gliovis.bioinfo.cnio.es/; Broad Institute Single Cell Portal (https://singlecell.broadinstitute.org/single_cell). See Materials and Methods for references. Data from this study are freely available for non-commercial purposes upon request to the corresponding author.

## Notes

### Competing Interest Statement

The authors have declared no competing interest.

