## Supplementary material for "Transglutaminase 2 function in glioblastoma tumor efferocytosis": Lui et al 2024 Supplemental

**SUPPLEMENTARY FIGURES**

*Figure S1. Mouse xenograft model of mesenchymal subtype glioblastoma.* PriGO17A cells were injected intracerebrally into CD1 nude mice. Mice were euthanized when they showed signs of morbidity and brains were formalin-fixed and paraffin-embedded. Immunohistochemistry for the macrophage/microglia marker Iba1 was performed on sections. A. Iba1 staining at site distant from tumour. Inset shows complete section with close-up region indicated with the small black box. Scale bar is 300 μm; B. Iba1 staining at site proximal to the tumour; C. Close-up image from (A), showing microglia morphology; D. Close-up image from B, showing macrophage/microglia morphology proximal to tumour. Macrophage/microglia have darker Iba1 staining and a less ramified morphology, consistent with an activated phenotype; E. Expression of *CCL2* mRNA in PriGO17A cells relative to three classical subtype glioblastoma cell patient isolates determined by microarray expression analysis. D. Secreted CCL2 protein concentrations, normalized to cell numbers, for PriGO17A cells and three classical subtype glioblastoma cell patient isolates. Data show the mean and standard deviation of four replicates. * indicates $P < 0.001$ by One Way Analysis of Variance and All Pairwise Multiple Comparison Procedures (Holm-Sidak method) for 17A versus all other samples.

*Figure S2. TGM2 antibody validation by Western blotting.* A. THP-1 cells were either differentiated into macrophages with PMA or undifferentiated as described in Materials and Methods. Differentiated and undifferentiated cells were also treated further with either IL4 or IFNγ as indicated. Whole cell extracts were then collected and analyzed for TGM2 expression by Western blotting. Amido black staining and GAPDH antibody were used as loading controls. K562 whole cell extract was used as a positive control. B. PMA-differentiated THP-1 cells were treated with TGFβ for the indicated times. Whole cell lysates were then analyzed for TGM2 expression as in A. C. PriGO17A glioblastoma cells were treated as indicated and whole cell lysates were then analyzed for TGM2 expression; D. Control staining with non-specific rabbit IgG. Immunohistochemistry was performed on mouse xenograft sections using rabbit non-specific monoclonal IgG control at the same concentration that was used for the TGM2-specific rabbit monoclonal. Top panel show an image from the un-injected hemisphere while the bottom panel shows an image from the injected hemisphere.

*Figure S3. Double immunofluorescence with Iba1 and TGM2 antibodies on xenograft sections.* Top panels show Iba1 (red), TGM2 (green) and merged image. Scale bar is 100 μm. The smaller and larger white boxes indicate the regions for the close-up images in the middle and bottom panels, respectively. Middle panels show close-up images of two microglia/macrophages from the images in the top panels. Bottom panels show close-up images of a necrotic region from the top panels.

*Figure S4. Examples of TGM2 immunohistochemistry on tissue microarray samples and TUNEL assays in patient organoids.* A. Example of TGM2 staining in tumour vasculature from TMA; B. Example of sporadic TGM2 positive cells within the tumour mass, similar to what was observed in the xenograft model; C. Image of a whole TMA core showing a region with cells that stain either moderately or strongly for TGM2; D. Close-up shows a strong TGM2-positive cell engaged in efferocytosis by morphological criteria (indicated with red arrow). E. Iba1 immunofluorescence (green) and TUNEL assay (red) on Patient C organoid samples that were untreated (control) or treated with either ONO-7475 for 48 h or NC9 for 24 h. The control sample is the same as shown in Figure 6B, with the DAPI signal removed to show the overall Iba1-positive cell interactions with apoptotic cells more clearly.

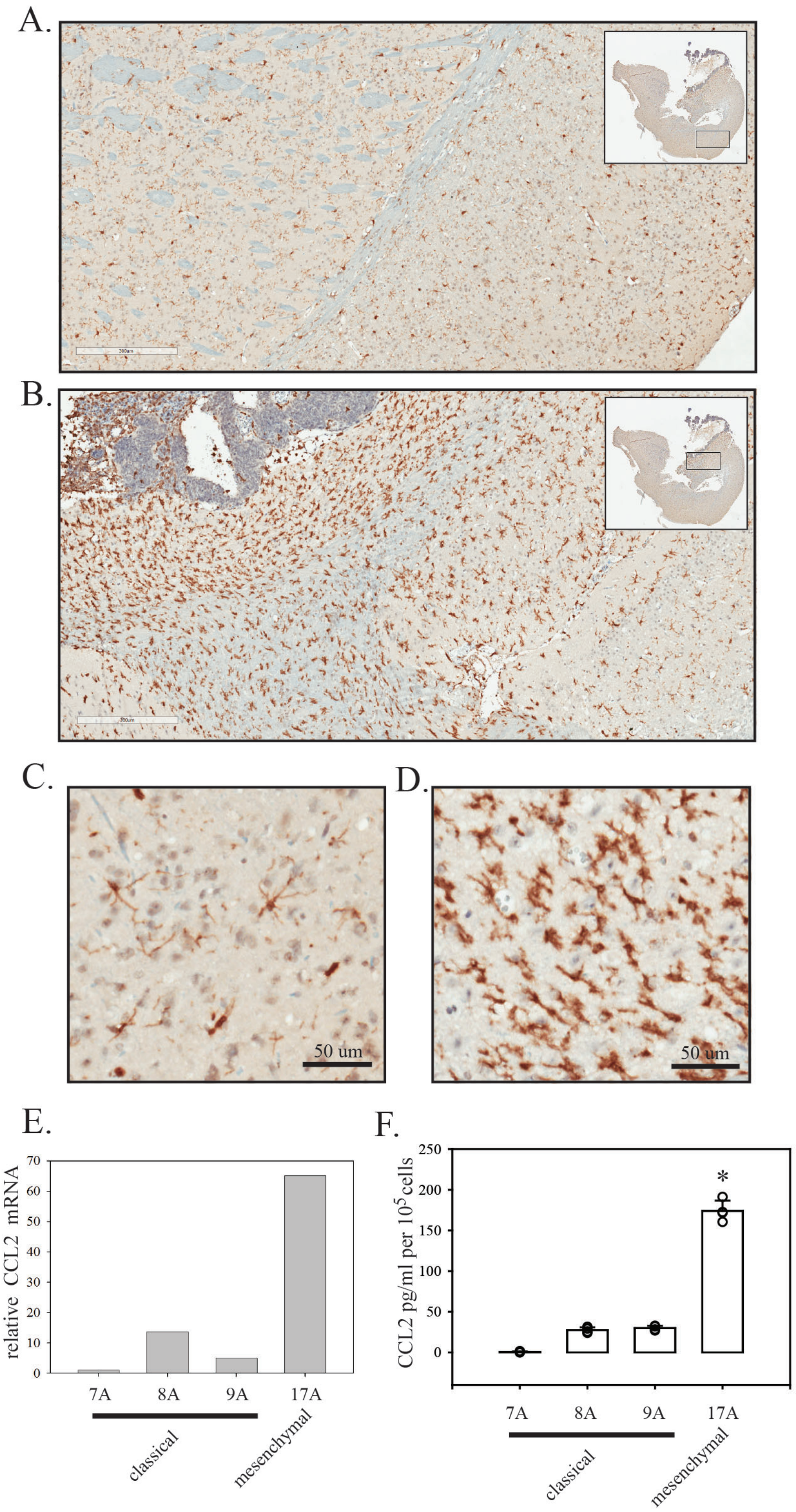
A.
B.
C.
D.
50 um
50 um
E.
relative CCL2 mRNA
70
60
50
40
30
20
10
0
7A
8A
9A
17A
classical
mesenchymal
F.
CCL2 pg/ml per 10$^5$ cells
250
200
150
100
50
0
*
7A
8A
9A
17A
classical
mesenchymal

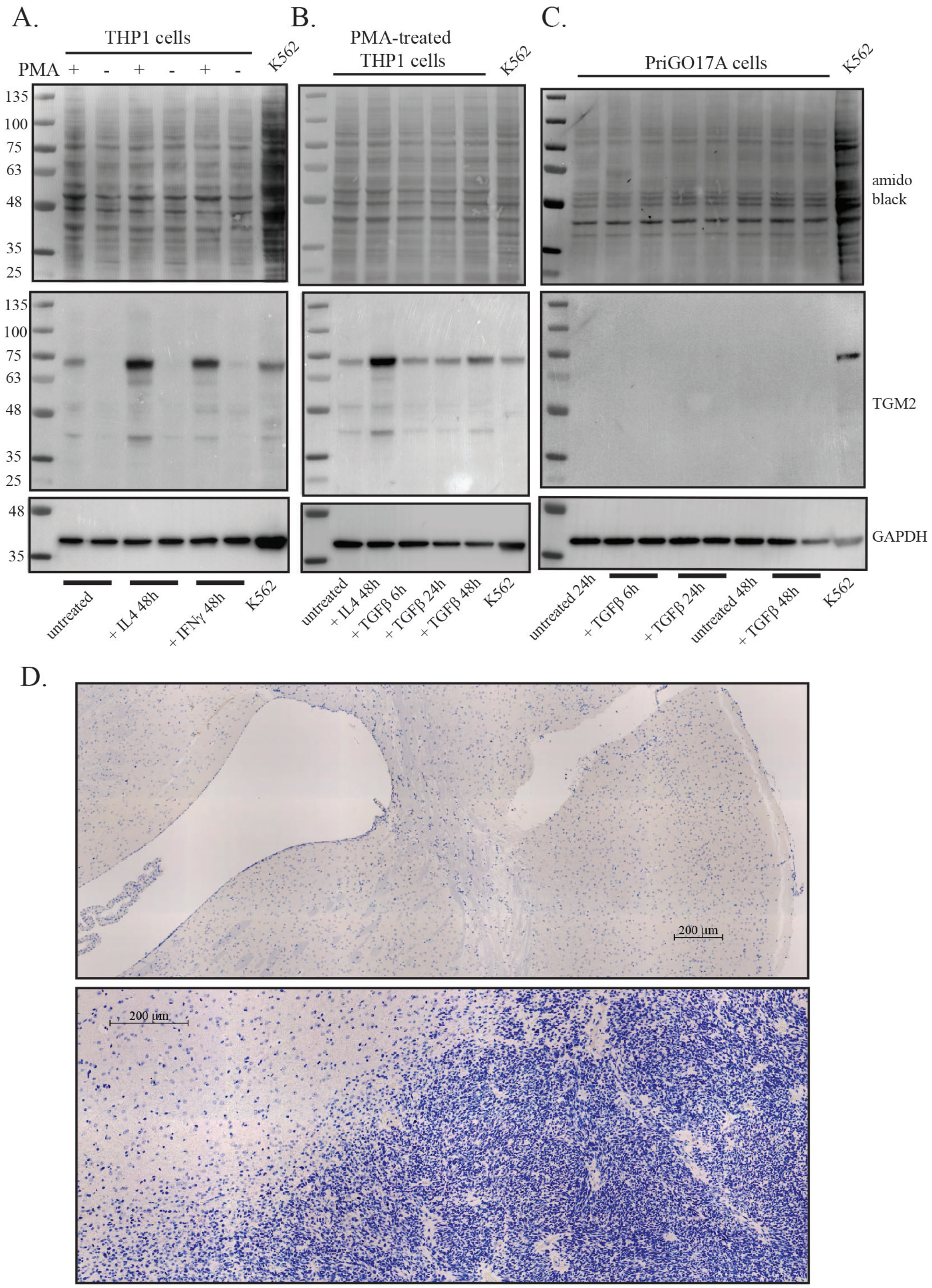

A.
THP1 cells
PMA + - + - + -
K562
B.
PMA-treated THP1 cells
K562
C.
PriGO17A cells
K562
135
100
75
63
48
35
25
amido black
TGM2
GAPDH
untreated
+ IL4 48h
+ IFNγ 48h
K562
untreated
+ IL4 48h
+ TGFβ 6h
+ TGFβ 24h
+ TGFβ 48h
K562
untreated 24h
+ TGFβ 6h
+ TGFβ 24h
untreated 48h
+ TGFβ 48h
K562
D.
200 µm
200 µm

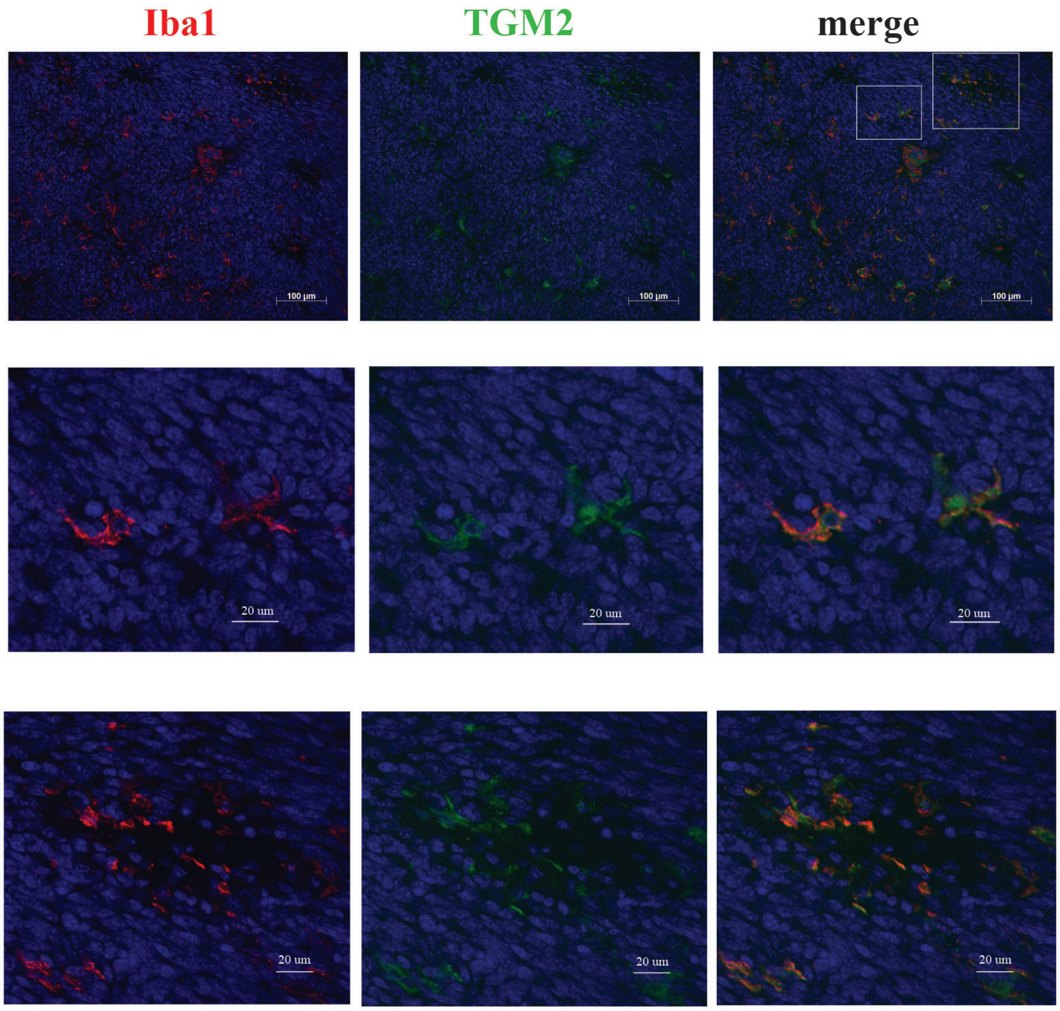
Iba1
TGM2
merge
100 µm
100 µm
100 µm
20 um
20 um
20 um
20 um
20 um
20 um

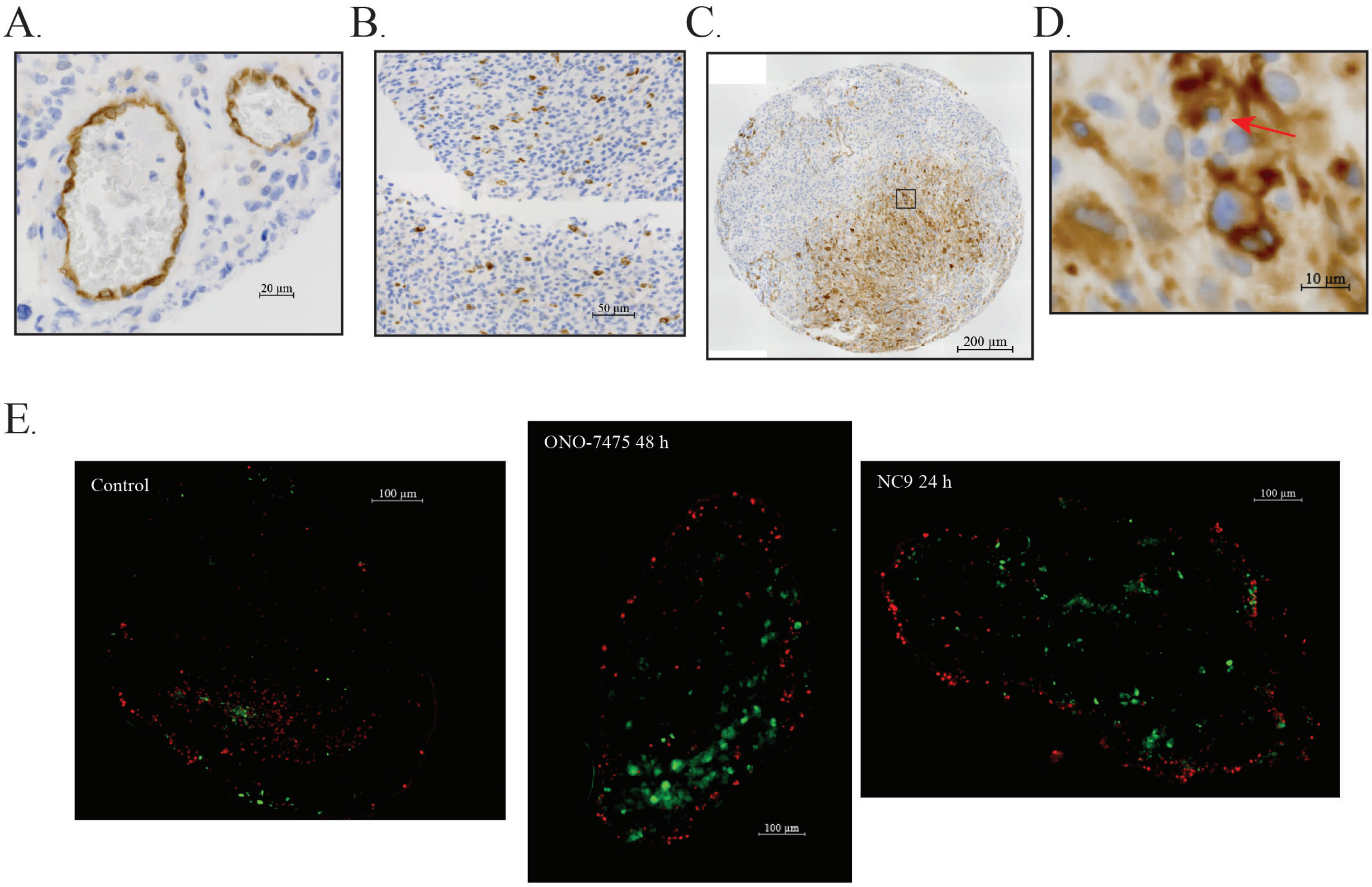

A.
20 µm
B.
50 µm
C.
200 µm
D.
10 µm
E.
Control
100 µm
ONO-7475 48 h
100 µm
NC9 24 h
100 µm
